# Single-cell transcriptomic analysis of the adult mouse spinal cord

**DOI:** 10.1101/2020.03.16.992958

**Authors:** Jacob A. Blum, Sandy Klemm, Lisa Nakayama, Arwa Kathiria, Kevin A. Guttenplan, Phuong T. Hoang, Jennifer L. Shadrach, Julia A. Kaltschmidt, William J. Greenleaf, Aaron D. Gitler

**Affiliations:** Department of Genetics, Stanford University School of Medicine, Stanford, CA 94305, USA; Stanford Neurosciences Graduate Program, Stanford University School of Medicine, Stanford, CA 94305, USA; Department of Neurosurgery, Stanford University School of Medicine, Stanford, CA 94305, USA; Department of Neurology and Neurological Science, Stanford University School of Medicine, Stanford, CA 94305, USA; Department of Applied Physics, Stanford University, Stanford, CA 94305, USA; Chan Zuckerberg Biohub, San Francisco, CA, USA

## Abstract

The spinal cord is a fascinating structure responsible for coordinating all movement in vertebrates. Spinal motor neurons control the activity of virtually every organ and muscle throughout the body by transmitting signals that originate in the spinal cord. These neurons are remarkably heterogeneous in their activity and innervation targets. However, because motor neurons represent only a small fraction of cells within the spinal cord and are difficult to isolate, the full complement of motor neuron subtypes remains unknown. Here we comprehensively describe the molecular heterogeneity of motor neurons within the adult spinal cord. We profiled 43,890 single-nucleus transcriptomes using fluorescence-activated nuclei sorting to enrich for spinal motor neuron nuclei. These data reveal a transcriptional map of the adult mammalian spinal cord and the first unbiased characterization of all transcriptionally distinct autonomic and somatic spinal motor neuron subpopulations. We identify 16 sympathetic motor neuron subtypes that segregate spatially along the spinal cord. Many of these subtypes selectively express specific hormones and receptors, suggesting neuromodulatory signaling within the autonomic nervous system. We describe skeletal motor neuron heterogeneity in the adult spinal cord, revealing numerous novel markers that distinguish alpha and gamma motor neurons—cell populations that are specifically affected in neurodegenerative disease. We also provide evidence for a novel transcriptional subpopulation of skeletal motor neurons. Collectively, these data provide a single-cell transcriptional atlas for investigating motor neuron diversity as well as the cellular and molecular basis of motor neuron function in health and disease.

## Main

The central nervous system (CNS) receives sensory input from its surroundings, integrates that information, and then communicates with muscles and organs throughout the body to enact an appropriate response. While a vast network of interconnected neurons is responsible for processing information and planning motor behaviors^1,2^, the transmission of signals from the CNS to peripheral tissues is controlled by one special and exceptionally rare cell population: spinal motor neurons.

Spinal motor neurons are unique because they reside in the CNS yet extend their axons far into the periphery in order to reach their innervation targets. Their activity is essential for virtually all skeletal and smooth muscle contractions in the body, controlling actions that range from the regulation of blood pressure^3^ and sweat secretion^4^ to the contraction of fast-twitch muscle fibers^5^. Because spinal motor neurons play an integral role in such diverse systems within the body, we hypothesized that they would have similarly heterogeneous gene expression profiles.

Considering their physiological importance and stoichiometric rarity, it is not surprising that spinal motor neuron dysfunction underlies many human neuromuscular diseases. Indeed, spinal motor neuron degeneration is causally responsible for amyotrophic lateral sclerosis (ALS), spinal muscular atrophy (SMA), and numerous other rare neuromuscular disorders^6^. In many of these diseases, certain populations of spinal motor neurons are uniquely affected, while others are spared^7,8^. Defining the transcriptional differences that separate these susceptible and resistant populations could lead to a better understanding of the molecular underpinnings of susceptibility, thus opening novel avenues for developing targeted interventions.

Despite their critical functional role, spinal motor neurons make up less than 0.4% of the total cells in the mammalian spinal cord^9^. This rarity has made them notoriously difficult to transcriptionally characterize^10^. To overcome these hurdles, we developed a motor neuron enrichment strategy using a fluorescent reporter mouse. This yielded an ∼100-fold increase in representation of motor neuron nuclei over past efforts^11^, enabling us to investigate how distinct transcriptional programs underlie spinal motor neuron function.

We performed single-nucleus RNA sequencing on 43,890 nuclei from the adult mouse spinal cord, providing us with unprecedented single-cell resolution of the spinal motor system. Here, we reveal numerous newly discovered marker genes to transcriptionally distinguish spinal motor neurons of the autonomic (visceral motor neurons) and somatic (skeletal motor neurons) nervous system. Furthermore, we provide evidence for previously unappreciated heterogeneity within both the somatic and autonomic nervous systems, including uncharacterized spinal motor neuron subtypes. These transcriptomes give key insights into the link between the brain and the body by defining the neuropeptides, transmitters, and receptors that motor neurons use to communicate. Furthermore, this detailed characterization will enable the development of transgenic mice and molecular tools that unlock genetic access to previously uncharacterized spinal motor neuron populations.

### Single-nucleus profiling of the adult mouse spinal cord

Because spinal motor neurons are so scarce, we enriched for motor neuron nuclei using a transgenic fluorescent reporter mouse^12^. This technique enabled us to selectively isolate cholinergic nuclei (*Chat+*), a population that encompasses all motor neurons and few other cell types in the adult mouse spinal cord^13^ (see Methods). Given the important role of cell non-autonomous mechanisms in neurodegeneration^14–16^, we also isolated non-motor neuron cells – including interneurons, astrocytes, microglia, and oligodendrocytes. In total, we transcriptionally profiled 43,890 nuclei from wild-type adult mice, mixing 20-40% of cholinergic nuclei with 60-80% from other cells in the spinal cord (Fig. 1a). We used graph-based methods to cluster nuclei and then annotate cell types according to averaged expression across clusters of common marker genes and genes encoding neurotransmitter signaling machinery (Fig. 1b, Methods). This approach enabled us to assign all captured nuclei into seven distinct categories: excitatory interneurons, inhibitory interneurons, cholinergic neurons, astrocytes, microglia, oligodendrocytes, and endothelial cells (Fig. 1c, Extended Data Fig. 1a-b). Based on these categories, we estimate that ∼30% (13,589 cells) of the profiled single-nucleus transcriptomes correspond to cholinergic neurons. This is a massive improvement in representation over prior efforts (∼50-100 motor neurons recovered)^11^ and provides unparalleled access to the transcriptional heterogeneity of spinal motor neurons.

**Figure 1:**
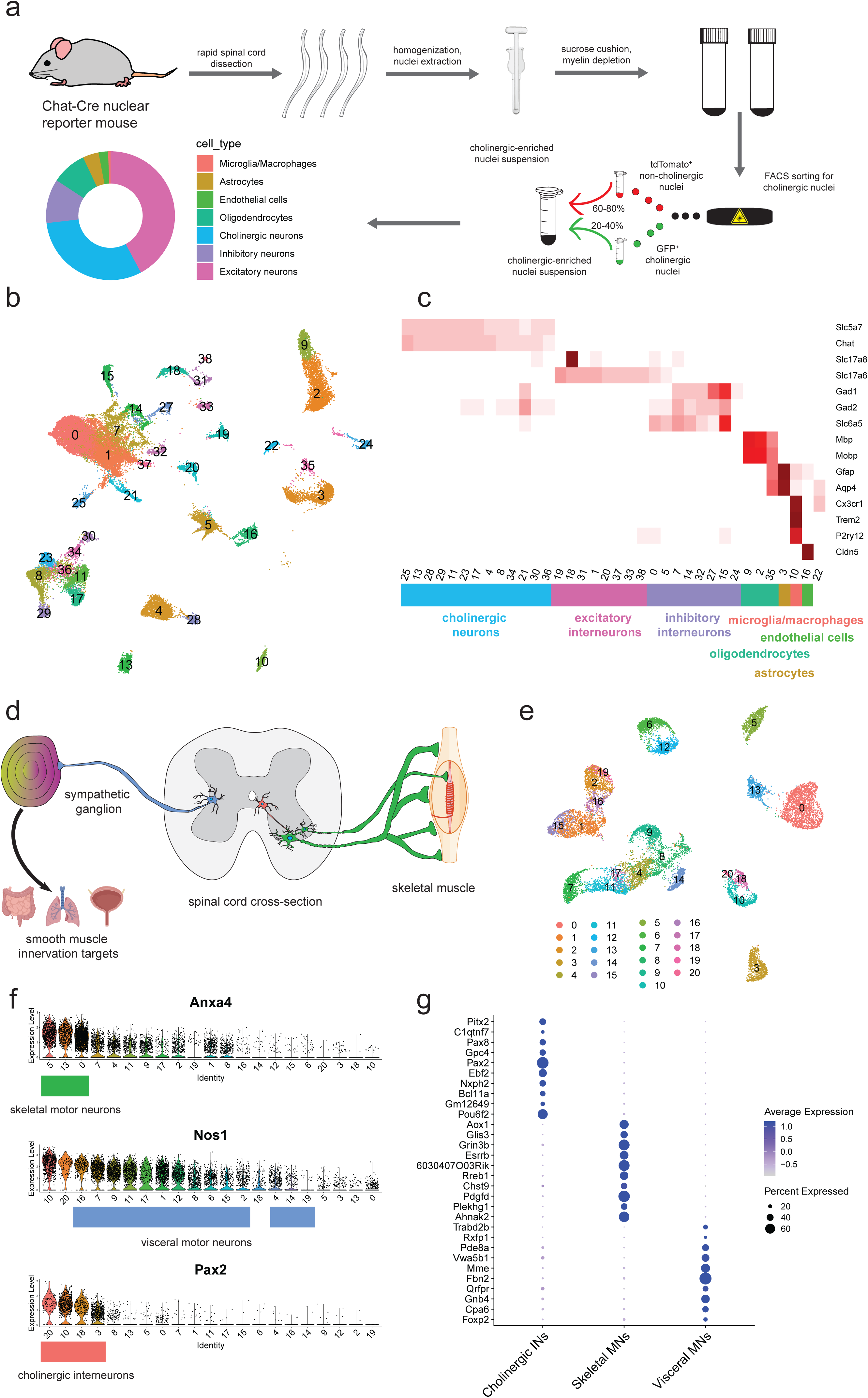
Motor neuron enrichment and single-nucleus transcriptional analysis of the adult mouse spinal cord uncovers novel skeletal and visceral motor neuron markers. **a**, Workflow for cholinergic nucleus enrichment and single-nucleus RNA sequencing (snRNAseq)—GFP+ and tdTomato+ cells were mixed at a ratio between 1:3 and 2:3. Plot shows the distribution of canonical cell types. **b**, UMAP of clustered snRNAseq data from 43,890 transcriptomes. **c**, Average expression levels per cluster for marker genes of each canonical cell population. Cell type labels based on expression patterns of marker genes. **d**, Schematic depicting expected cholinergic cell types in the spinal cord. Visceral motor neurons (blue) innervate sympathetic ganglia, skeletal motor neurons (green) directly innervate muscle fibers, and cholinergic interneurons (red) innervate motor neurons and other cells. **e**, UMAP with graph-based clustering of all cholinergic neurons reveals 21 clusters. **f**, Ranked expression of known marker genes *Pax2* (interneurons) and *Nos1* (visceral motor neurons), as well as novel marker gene *Anxa4* (skeletal motor neurons) by cluster. Cell labels were assigned hierarchically by expression levels of *Pax2, Nos1*, and finally *Anxa4* and are reported below each plot. **g**, Novel marker genes for cholinergic interneurons, skeletal motor neurons, and visceral motor neurons. Dot size is proportional to the percent of each cluster expressing the marker gene, while blue color intensity is correlated with expression level. All expression values were log-normalized in Seurat^72^.

### Novel genetic markers distinguish autonomic vs. skeletal motor neurons

We next asked whether the observed transcriptional diversity of spinal motor neurons corresponds to functionally defined cell types (Fig. 1d). We computationally isolated and used graph-based clustering (see Methods) to segregate all cholinergic neurons into 21 clusters (Fig. 1e). We annotated these subpopulations as skeletal motor neurons, cholinergic interneurons, and visceral motor neurons (Fig. 1f) based on expression of known marker genes as well as expression patterns of uncharacterized genes in the publicly available Allen Mouse Spinal Cord Atlas^17^. Specifically, we identified cholinergic interneurons based on their expression of *Pax2*^18^, while visceral motor neurons (which are part of the autonomic nervous system) were neuronal nitric oxide synthase positive (*Nos1+*), a characteristic feature of the autonomic nervous system^19–21^, and expressed high levels of *Fbn2* and *Zeb2* (Fig. 1g, Extended Data Fig. 1c). We confirmed that *Fbn2* and *Zeb2* were expressed specifically in the lateral autonomic columns of the thoracic and sacral spinal cord— where visceral motor neurons are specifically located (Extended Data Fig. 2a-b).

Because there are no known robust skeletal motor neuron markers, we hypothesized that the remaining three clusters represent skeletal motor neurons. To test this hypothesis, we examined the Allen Mouse Spinal Cord Atlas for expression of several of the top marker genes from those clusters—such as *Tns1* and *Anxa4*. Indeed, both genes are strongly expressed in small and large diameter neurons in the ventral horn of the spinal cord—a pattern that is consistent with skeletal motor neurons (Extended Data Fig. 2c-b)^22^. Our classification of skeletal and visceral motor neurons differs from a previous study, which postulated that both *Fbn2* and *Zeb2* expression were novel markers specifically expressed in alpha (α) motor neurons^23^. Instead, we find that *Zeb2* is almost entirely absent from skeletal motor neurons (Extended Data Fig. 2e)—an observation that is consistent with its role as a transcription factor that defines the visceral motor neuron lineage during development^20^. These new transcriptional profiles reveal dozens of new marker genes that may be used to more reliably distinguish skeletal motor neurons from other cells in the spinal cord. (Fig. 1g; Supp. Table S1a).

### Single-nucleus transcriptomics reveals diversity within the autonomic nervous system

In contrast to skeletal motor neurons, which control voluntary movement, visceral motor neurons in the spinal cord control the activity of involuntary smooth muscles responsible for regulating many homeostatic processes throughout the body, such as blood pressure and respiratory air flow^24,25^. These cells are part of the sympathetic nervous system, while comparable cells in the brain stem are part of the parasympathetic nervous system^25^. These sympathetic visceral motor neurons are developmentally very closely related to skeletal motor neurons^26^, but do not innervate muscle fibers directly. Instead, they predominantly project from the lateral autonomic column of the spinal cord and synapse onto the sympathetic chain ganglia that control smooth muscle contraction (Fig. 2a)^24^. Neurons of the autonomic nervous system innervate nearly every organ in the body and thus have different functional requirements to fit the needs of each organ (Fig. 2b). Could these differences be encoded by transcriptional heterogeneity?

**Figure 2:**
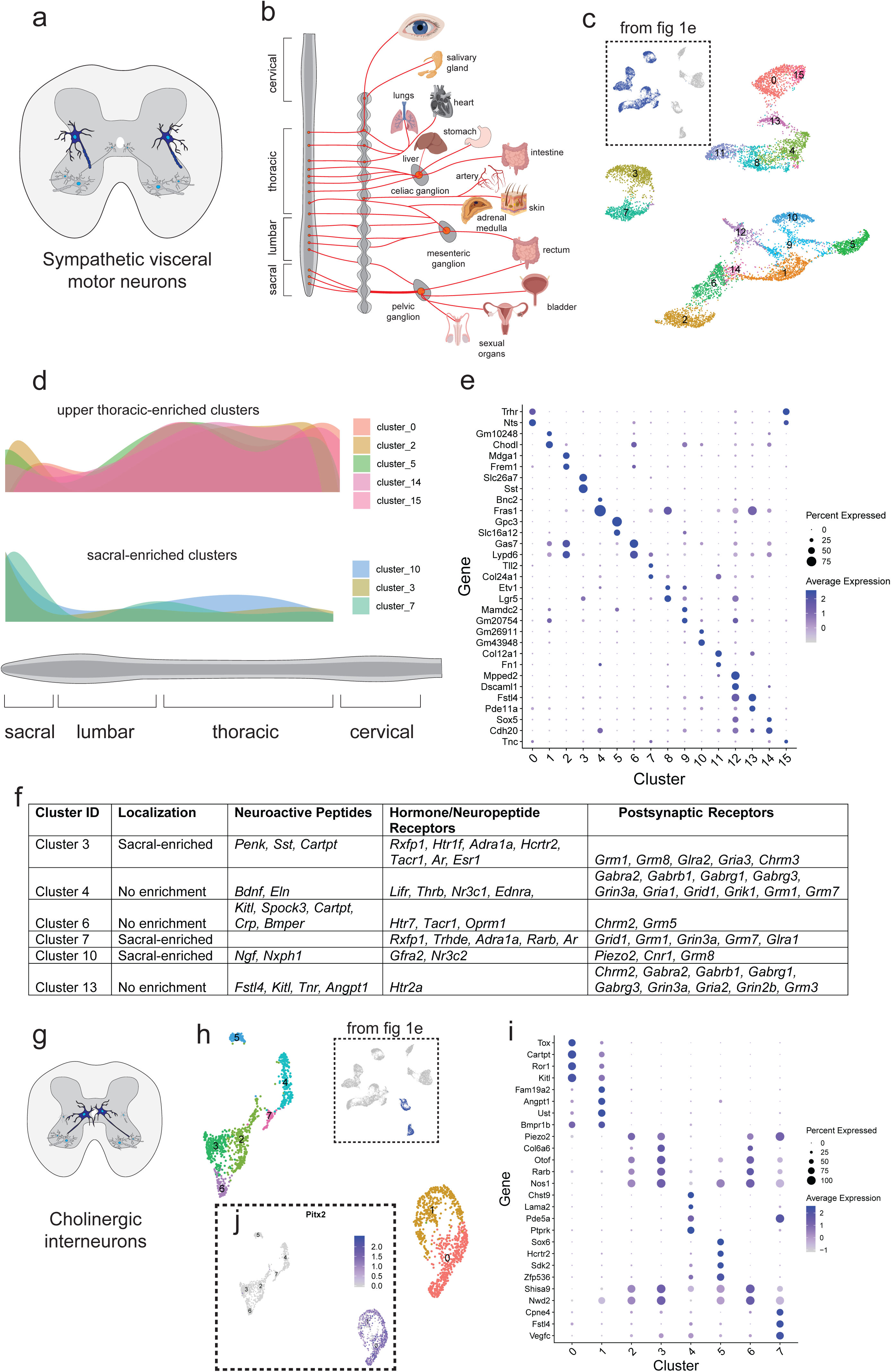
Single-nucleus transcriptomics reveals immense diversity within the autonomic nervous system and partition cells. **a**, Schematic illustrating the position of sympathetic visceral motor neurons (blue) in the lateral autonomic column of the spinal cord. **b**, Diagram adapted from Espinosa-Medina 2016 showing innervation targets of the sympathetic nervous system^25^. **c**, UMAP with 16 visceral motor neuron subclusters. Inset shows all cells from Fig 1e that were subclustered. **d**, Estimated relative density of visceral motor neurons along the rostral-caudal axis of the spinal cord. Density functions are combined density estimates of marker genes for each cluster (see Methods). Clusters were grouped according to shape of density function. **e**, Novel marker genes for each cluster of visceral motor neurons. **f**, Table highlighting marker gene expression and localization of several highlighted clusters of visceral motor neurons. Neuropeptides, hormone receptors, and neurotransmitter receptors were annotated with DAVID gene ontology. All genes listed show differential enrichment in one cluster compared with all others (DESeq2, p_adj<0.01, log2fc>0.5). **g**, Schematic showing cholinergic interneuron innervation of skeletal motor neurons as demonstrated previously^34^. **h**, UMAP with graph-based clustering labels from subclustering all cholinergic interneurons. Inset shows all cells from Fig1e that were subclustered. **i**, Novel marker genes for cholinergic interneuron clusters identified. **j**, Expression of *Pitx2* in cholinergic interneuron populations, overlaid on UMAP projection from (h). Cluster 0 and 1 are *Pitx2*-positive. All expression values were log-normalized in Seurat^72^. Dot size is proportional to the percent of each cluster expressing the marker gene, while blue color intensity is correlated with expression level in **(e**,**i)**.

To test this hypothesis, we subclustered all visceral motor neuron transcriptomes and found 16 transcriptionally distinct populations (Fig. 2c). The sympathetic nervous system is known to be organized spatially along the rostral-caudal axis of the spinal cord, such that visceral motor neuron innervation targets localize to specific spinal cord levels (Fig. 2b)^24^. We hypothesized that transcriptionally distinct clusters reflect different anatomic positions and/or innervation targets. We identified genes with expression patterns in the adult mouse lateral autonomic column of the spinal cord using *in situ* hybridizations in the Allen Spinal Cord Atlas^17^. From this list, we selected genes whose expression varied across the 16 clusters we identified. Then, for each gene we counted the number of cells expressing it in various levels of the spinal cord, and fit polynomial density distributions to these observations (see Methods). We performed weighted averaging of these distributions based on relative expression levels of the marker genes from each cluster, yielding an estimate of the positional distribution of cells within each cluster along the rostral-caudal spinal cord axis (Fig. 2d) compared with the underlying distribution (Extended Data Fig. 3a, Methods). While some clusters showed no bias along this axis (Extended Data Fig. 3a), several showed a clear enrichment in the sacral (clusters 3, 7, 10, Fig. 2d) and upper-thoracic (clusters 0,2,5,14,15; Fig. 2d) regions of the spinal cord.

We found preganglionic clusters to be transcriptionally highly divergent — with numerous specific markers present in most clusters (Fig. 2e). Because it is well known that neuropeptides play a crucial role in the sympathetic nervous system^27,28^, we were specifically interested in differentially expressed genes that would affect those pathways. We observe remarkable specificity of neuropeptides and receptors—as well as neurotransmitter receptors—across visceral motor neuron clusters (Fig. 2f). Cluster 3, which is primarily located in the sacral spinal cord, expresses somatostatin (*Sst*) and proenkephalin (*Penk*) (Extended Data Fig. 3b-b). Each of these peptides has been shown to have neuromodulatory effects in the sympathetic ganglia^29–31^, suggesting that preganglionic motor activity may modulate post-ganglionic motor neuron activity. We also find highly specific expression of neuroactive peptide precursors in other clusters, indicating that neuropeptide expression may be a defining feature of visceral motor neuron populations (Supp. Table S1b).

Strikingly, visceral motor neuron subpopulations also express unique sets of hormone receptors known to modulate neuronal activity^32^. For instance, the relaxin family peptide receptor (*Rxfp1*) is specifically expressed in clusters 3 and 7 (Fig. 2f, Extended Data Fig. 3c), as are many others that involve serotonergic, adrenergic, cannabinoid, and opioid signaling (Fig. 2f). Transcriptional differences also offer key insights into pre- and post-synaptic organization. Several clusters are enriched for expression of GABA receptors *Gabra1, Gabrg1, Gabrb1, Gabrg3* (clusters 4 and 13, Fig. 2f), while others preferentially express glycine receptors *Glra1* and *Glra2* (clusters 7 and 3, respectively; Fig. 2f). These characteristics of unique autonomic populations will enable more targeted study—and manipulation—of populations that control highly divergent smooth muscle pools throughout the body.

### Transcriptional characterization of cholinergic interneurons

Cholinergic interneurons are a rare cell population marked by *Pax2* expression^18^. They play key roles in the circuits underlying locomotor behaviors^33^ (Fig. 2g). Subclustering revealed 7 distinct transcriptional populations of cholinergic interneurons (Fig. 2h) —including clusters 2,3,5,6, and 7 that express high levels of *Nos1*, which is traditionally considered to be a marker of the autonomic nervous system in the spinal cord^20,21^ (Fig. 2i). It remains to be determined whether this is a new population of *Nos1*-positive interneurons, or instead a preganglionic motor neuron population that projects into the periphery but also expresses the interneuron marker *Pax2*.

In contrast, clusters 0 and 1 do not express *Nos1* but instead express *Pitx2*—an established marker of partition cells^34,35^ (Fig. 2j). Partition cells are a subset of cholinergic interneurons that make direct cholinergic synapses with motor neuron somata and proximal dendrites. These synapses, referred to as ‘C boutons,’ modulate motor neuron excitability during locomotor activity^34,36^. Elegant viral tracing studies have further delineated partition cells into ipsilaterally and contralaterally projecting populations that make exquisitely specific synaptic connections with motor neurons^37^. We find a parallel transcriptional bifurcation in partition cells, suggesting that clusters 0 and 1 correspond to previously identified subtypes of partition cells.

By identifying partition cells using known markers, we were able to ask which genes they specifically express. Of particular interest is *Gldn*, a gene in our dataset whose expression is highly specific to partition cells among all spinal cord populations (Supp. Table S1a). Mutations in *GLDN* cause lethal congenital contracture syndrome, a crippling neurodegenerative disease in which joints become permanently fixed in a bent or straight position^38^. We suggest that mutations in *GLDN* could impair partition cell function, disrupting locomotor circuits early in development and possibly contributing to disease.

### Identification of novel alpha and gamma motor neuron markers

Skeletal motor neurons have traditionally been defined based on their muscle innervation target^4,39–41^, developmental lineage^26,42,43^, morphology^22,44^, and electrophysiological properties^45–48^. Accordingly, they are broadly classified as alpha (α), beta (β), and gamma (γ) spinal motor neurons (Fig. 3a). α motor neurons directly innervate extrafusal muscle fiber neuromuscular junctions (NMJs). In contrast, γ MNs innervate intrafusal muscle spindles. We identified skeletal motor neurons by *Tns1/Anxa4* expression (as above, see Methods), and then subclustered them (Fig. 3b).

**Figure 3:**
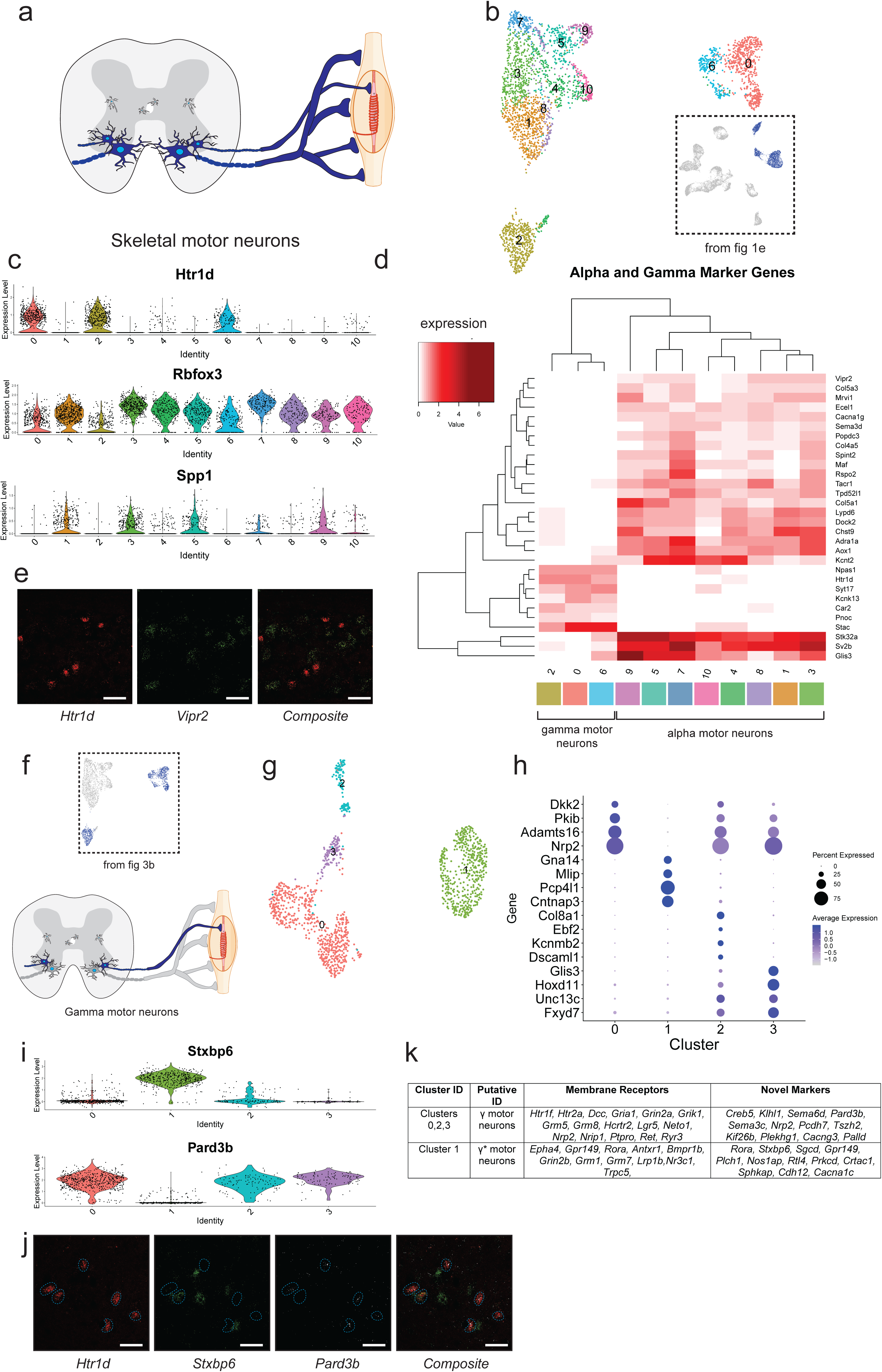
Transcriptional differences between alpha (α) and gamma (γ) motor neurons. **a**, Transverse schematic illustrating position of skeletal motor neurons (blue) in the ventral horn of the spinal cord. Gamma motor neurons are small and innervate intrafusal muscle fibers. α motor neurons are large and innervate extrafusal fibers. **b**, UMAP with 11 subclustered skeletal motor neurons populations. Inset shows all cells from Fig 1e that were subclustered. **c**, Average expression of known γ marker *Htr1d* and α markers *Rbfox3* and *Spp1* by cluster. **d**, Heatmap with average expression by cluster of differentially expressed genes in α and γ populations. Differentially expressed genes between γ and α populations. **e**, *In situ* hybridization against *Htr1d* and *Vipr2* in ventral horn of transverse spinal cord section. Composite demonstrates mutually exclusive expression. **f**, Transverse schematic illustrating γ motor neurons (blue) innervating intrafusal muscle fibers. Inset shows all cells from Fig 3b that were subsequently subclustered. **g**, UMAP with 4 populations of γ motor neurons. **h**, Novel marker gene expression across γ motor neuron subpopulations. Dot size is proportional to the percent of each cluster expressing the marker gene, while blue color intensity is correlated with expression level. **i**, *Stxbp6* and *Pard3b* bifurcate γ population. Average expression of *Stxbp6* and *Pard3b* gene across each cluster is shown. **j**, *In situ* hybridization against *Htr1d, Stxbp6*, and *Pard3b* in ventral horn of transverse spinal cord section shows reciprocal expression in subpopulations. White dotted lines mask *Htr1d*+ cells. **k**, Differentially expressed membrane receptors between two main populations of γ motor neurons, as well as novel markers that delineate them. All differential expression calculated using DESeq2 (p_adj < 0.01, log2fc>0.5). All expression values were log-normalized in Seurat^72^. Scale bars=50 µm.

To identify putative α and γ motor neuron populations, we examined marker gene expression within each cluster (Fig. 3c). There are few robust markers in adult animals^22,44,49^, but γ motor neurons express *Htr1d*^50^, while α motor neurons express high levels of *Rbfox3*^51^ and *Spp1*^52^. Clusters 0, 2, and 6 express high levels of *Htr1d* and low levels of *Rbfox3* and *Spp1* (Fig. 3c, Supp. Table S1c), suggesting that they represent γ motor neurons. The remaining clusters have low levels of *Htr1d* expression, suggesting that they represent α motor neurons (Fig. 3c). We calculated differential gene expression between putative α and γ motor neuron clusters, yielding a collection of novel markers of each population (Fig. 3d). To validate our markers, we performed *in situ* hybridization with a canonical γ marker (*Htr1d*) and a novel α marker (*Vipr2*), confirming their reciprocal expression pattern in cells of the ventral horn of the spinal cord (Fig. 3e). *Vipr2+* cells are larger than *Htr1d+* cells, recapitulating a distinguishing feature of α and γ motor neurons^49^. While all existing α motor neuron markers are insufficient on their own to distinguish α motor neurons from other cells in the spinal cord. *Vipr2* is expressed robustly and exclusively in all α motor neuron populations^22^ (Extended Data Fig. 4a). Together, these results provide a robust, novel molecular basis for distinguishing α and γ motor neurons.

### Transcriptional profiles of γ spinal motor neurons reveal two highly divergent motor neuron types

Gamma motor neurons innervate intrafusal muscle fibers, which maintain the tension required for skeletal muscle to function properly (Fig. 3f). Subclustering of γ motor neurons (*Htr1d*+) revealed four clusters (Fig. 3g) with numerous differentially expressed transcripts among them, such as *Hoxd11* in cluster 2 (Fig. 3h). *Hoxd11* is a transcription factor expressed in lumbar spinal motor neurons during development^53^, suggesting that some of the smaller clusters may reflect spatial position along the rostral-caudal axis of the spinal cord. We next used these top marker genes to hierarchically cluster all γ motor neurons and were surprised to find a dramatic divide between two subpopulations of γ motor neurons (Extended Data Fig. 4b, Supp. Table S1d). Curiously, many of the genes enriched in one of these populations are also expressed by α motor neurons (Extended Data Fig. 4c). We named this new population γ * (cluster 1), while all other cells are γ motor neurons (clusters 0,2,3). We can reliably distinguish γ from γ * solely by reciprocal expression of *Stxbp6* (γ *) or *Pard3b* (γ), both in our single cell dataset (Fig. 3i) and by *in situ* hybridization (Fig. 3j). These distinct populations likely represent a fundamental subdivision of the fusimotor system.

There are several skeletal motor neuron populations that have been historically defined physiologically or anatomically but not yet transcriptionally. Among these are α motor neurons– cells that innervate both intrafusal and extrafusal fibers—and therefore have properties of both α and γ motor neurons^54,55^. Could γ* actually correspond to this historically elusive and long-sought skeletal motor neuron subtype? We present a list of novel markers that differentiate γ and γ* populations (Fig. 3k), which will enable a more detailed exploration of this hypothesis.

### Transcriptional analysis of α motor neurons reveals distinct pools

α motor neurons innervate extrafusal muscle fibers^22^ (Fig. 4a). Owing to a vast heterogeneity in muscle location throughout the body and the types of muscle contractions required for coordinated movement, α motor neurons have substantial functional differences^56–58^. To define differences at the transcriptional level, we subclustered the α motor neuron transcriptomes (as above—see Methods; Fig. 4b). This analysis revealed 12 clusters. Most α motor neurons fall in one large population that consists of clusters 0,1, and 4. Other clusters are transcriptionally distinct from this main population, and expression of markers in these populations reveals a high degree of specificity (Fig. 4c). Intriguingly, cluster 3 highly expresses *Cpne4* and *Fign* (Fig. 4c), which were recently shown to be highly expressed in digit-innervating motor neurons^56^. Furthermore, clusters 8 and 9 express high levels of *Sema3e* (Fig. 4c)—which is a genetic marker for shoulder-innervating motor neurons in the cervical spinal cord. *Sema3e* encodes a protein that, together with its receptor PlexinD1^59,60^, is responsible for synaptic specificity between sensory and motor neurons in that pool^61^. These findings raise the hypothesis that transcriptional subpopulations of α motor neurons correspond to motor pools that specifically innervate distinct muscle groups.

**Figure 4:**
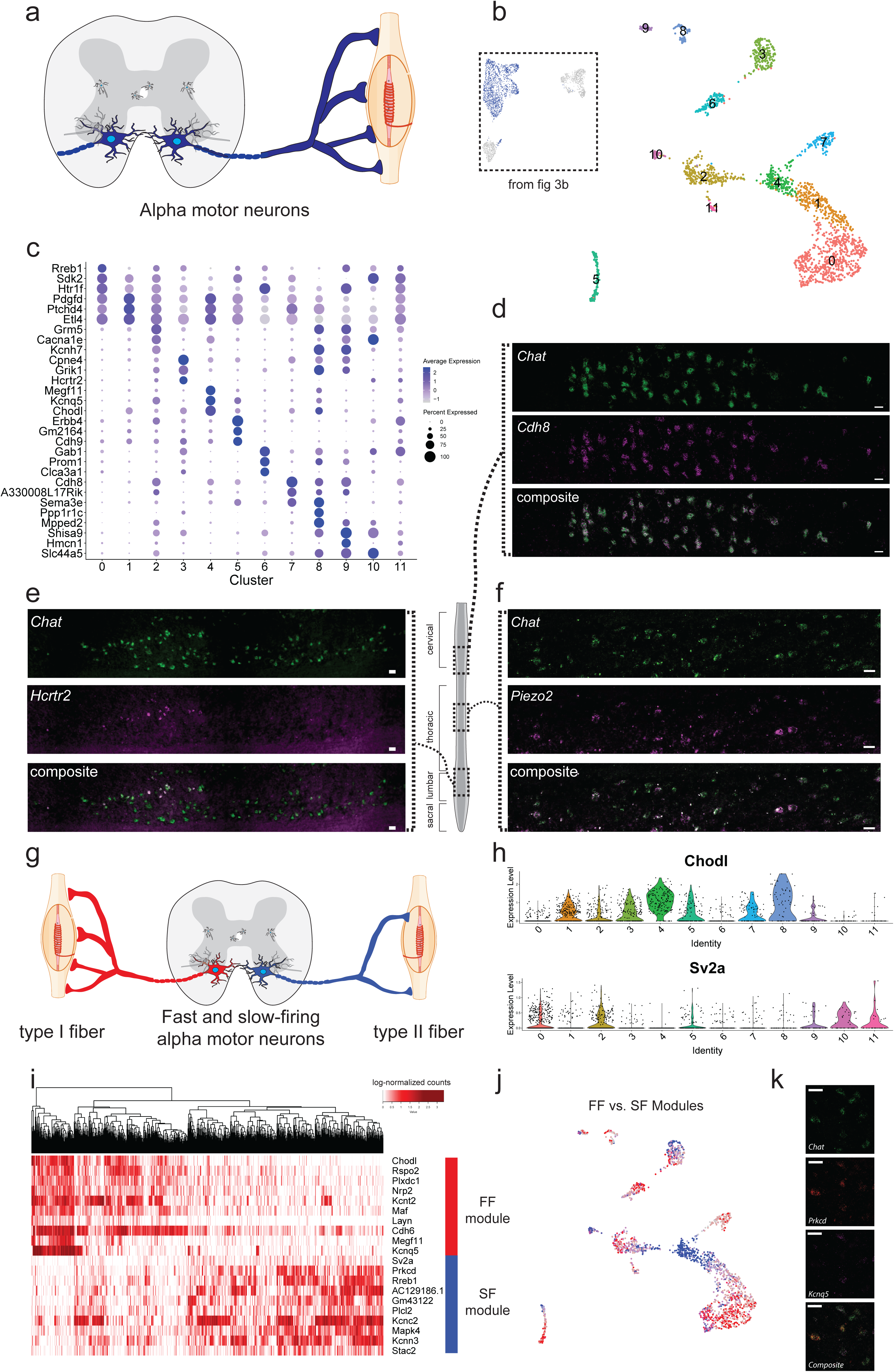
Alpha (α) motor neuron pool, position, and homeostatic properties reflect transcriptional differences. **a**, Transverse schematic shows α motor neurons (blue) innervating extrafusal muscle fibers. **b**, UMAP with 12 subclustered α motor neuron populations. Inset shows all α motor neurons from Fig 3b were subclustered. **c**, Novel marker gene expression across α motor neuron subpopulations. Dot size is proportional to the percent of each cluster expressing the marker gene, while blue color intensity is correlated with expression level. **d**, *In situ* hybridization against *Chat* and *Cdh8* in sagittal spinal cord section. Overlay shows enrichment of double-positive cells in cervical spinal cord. Schematic shows position within the spinal cord where images were taken (full images in Fig S6a). **e**, *In situ* hybridization against *Chat* and *Hcrtr2* in sagittal spinal cord section. Overlay shows enrichment of double-positive cells in motor pool of lumbar spinal cord. Schematic shows position within the spinal cord where images were taken (full images in Fig S6b). **f**, *In situ* hybridization against *Chat* and *Piezo2* in sagittal spinal cord section. Overlay shows numerous double-positive cells in thoracic spinal cord. **g**, Schematic illustrating slow (red) and fast-firing (blue) α motor neuron populations innervating type I and type II fibers. **h**, Expression of known fast-firing (*Chodl*) and slow-firing (*Sv2a*) marker genes across all clusters. **i**, Heatmap with all α motor neurons, hierarchically clustered and colored by expression differentially expressed genes between FF and SF α motor neurons. Red and blue bars show gene modules enriched in FF and SF α motor neurons, respectively. **j**, Expression of FF and SF gene modules overlaid on UMAP of all α motor neurons. **k**, *In situ* hybridization against *Chat, Prkcd, and Kcnq5* in ventral horn of transverse spinal cord section. Overlay shows reciprocal expression of *Prkcd* and *Kcnq5* in *Chat+* cells. All differential expression calculated using DESeq2 (p_adj < 0.01, log2fc>0.5). All expression values were log-normalized in Seurat^72^. Scale bars=50 µm.

To test whether the α motor neuron clusters correspond to these previously characterized motor pools, we generated orthogonal markers of presumed digit-innervating (cluster 3) and shoulder muscle-innervating motor neurons that do not have known expression patterns in the spinal cord. *Cdh8*, whose expression is similar to *Sema3e* (Fig. 4c), localized to motor neurons in the cervical spinal cord—the same position where the shoulder-innervating motor pool would be expected^61^ (Fig. 4d, Extended Data Fig. 5b). *Hcrtr2*, a gene that encodes a membrane-bound hypocretin (orexin) receptor^62^ and is correlated with *Cpne4* expression in α motor neurons (Extended Data Fig. 5c), localized specifically to a pool of motor neurons in the lumbar and thoracic spinal cord (Fig. 4e, Extended Data Fig. 5d). This mirrors the spatial organization of the previously described digit-innervating motor neurons^56^. Collectively, these data provide evidence that transcriptional heterogeneity underlies motor pool identity and likely function in the adult mouse spinal cord. The abundance of specifically expressed genes in each novel population will empower further functional study of these motor pools (Supp. Table S1e).

Unexpectedly, we found *Piezo2* is highly expressed in α motor neurons (Fig. 4f). *Piezo2* is best known for its role in somatosensory neurons. It encodes a mechanosensitive cation channel that opens in response to physical displacement of the cell membrane^63,64^. We used *in situ* hybridization to confirm these findings—almost all α motor neurons show some level of expression, and interspersed are sporadic high expressors. Given the surprisingly high *Piezo2* expression in α motor neurons, we hypothesize that it may have a functional role at the neuromuscular junction. Conditional gene inactivation studies will help to define Piezo2’s functional role in motor neurons.

### Fast and slow-firing α motor neurons have divergent transcriptional signatures

Skeletal muscles innervated by motor neurons are composed of slow (type I) and fast (type II) twitch fibers, which exhibit different patterns of synaptic input (Fig. 4g)^65^. Likewise, α spinal motor neurons have different electrophysiological and metabolic properties^22^. These range from fast-firing (FF) to slow-firing (SF), and importantly these classes have major differences in susceptibility to degeneration in ALS^8,44^. Thus, determining the transcriptional differences between these classes of motor neurons may provide insight into this differential susceptibility^8,65^. Molecularly, FF and SF neurons are distinguished by mutually exclusive expression of *Chodl* and *Sv2a* respectively^35^. We confirmed this striking anticorrelated expression pattern between *Chodl* and *Sv2a* across all α motor neuron clusters (Fig. 4h).

We next analyzed transcription profiles for gene expression modules that might underlie the functional properties that separate FF and SF motor neurons. We identified differentially expressed genes between *Chodl*^+^ and *Sv2a*^*+*^ α motor neurons, and then hierarchically clustered cells based on expression of this gene set (Fig. 4i). These genes, composed of a module enriched in FF motor neurons and another enriched in SF motor neurons, are sufficient to distinguish FF from SF motor neurons. Furthermore, the FF and SF gene modules are expressed in reciprocal populations of α motor neurons across virtually all motor pools, but in different proportions (Fig. 4j).

Intriguingly, both the FF and SF modules contain genes that encode subunits of voltage-gated potassium channels (*Kcnt2* and *Kcnq5* in FF, *Kcnn3* in SF). Potassium channel subtypes play a vital role in determining the resting membrane potential and basal firing rate of neurons. The differential expression of potassium channel isoforms suggests a mechanism through which α motor neuron electrophysiological properties are established^65,66^. We find that SF neurons specifically express *Prkcd*, which encodes a protein with a fundamental role in determining how cells respond to oxidative stress and DNA damage^67,68^. Intriguingly, FF motor neurons do not express *Prkcd*, but instead express high levels of *Prkcb*, which encodes a key regulator of autophagy^69^. Dysregulation of each of these pathways is thought to be fundamental to ALS ontology and progression^7^. We performed *in situ* hybridization to confirm that *Prkcd* and *Kcnq5* are expressed in a mutually exclusive pattern throughout motor neuron populations (Fig. 4k). Collectively, these data reveal a rich transcription basis for the functional diversity of fast- and slow-firing motor neurons.

### Discussion

We report here a detailed molecular characterization of the adult mammalian motor system at single-cell resolution. Using a transgenic motor neuron enrichment strategy, we have simultaneously discovered new divisions within the autonomic and somatic motor systems, while greatly expanding the transcriptional characterization of these populations.

Within the autonomic motor system we discovered 16 subpopulations of sympathetic visceral motor neurons. Visceral motor neuron clusters expressed entirely different repertoires of neuromodulatory peptides with known activities in the peripheral ganglia that they innervate, such as somatostatin, proenkephalin, brain-derived neurotrophic factor, and nerve growth factor. This suggests that sympathetic efferents that innervate peripheral ganglia may affect neurotransmission more broadly at those sites in unconventional ways, potentially even modulating the activity of peripheral sensory neurons and postganglionic motor neurons^24^. Together, our findings inspire the possibility that specific visceral motor neuron populations may someday be selectively targeted with therapies to treat autonomic dysfunction in humans.

Within the somatic motor system, we present novel α and γ motor neuron markers in addition to a molecularly uncharacterized population of skeletal motor neurons (γ*), which shares the expected features of the elusive β motor neuron population. Furthermore, we find that transcriptional subpopulations of α motor neurons correspond to previously described, distinct motor pools^56,61^. We propose that other transcriptionally distinct α motor neuron populations in our dataset may similarly correspond to specialized motor pools, and we offer numerous novel markers to facilitate testing this hypothesis.

Our analysis also reveals gene modules that are selectively expressed in electrophysiologically and metabolically distinct populations of fast and slow-firing α motor neurons. These differentially expressed genes offer insight into how motor neuron subtypes establish unique biophysical properties that are specifically matched to the properties of their muscle innervation targets^5,65^. Indeed, our analysis revealed that fast and slow-firing α motor neurons express a divergent collection of potassium channel subunits, which are specifically required to tune the resting membrane potential and firing rate in neurons^66^. These expression profiles may also help suggest approaches to rescue aberrant function in fast-firing motor neurons, which specifically degenerate in ALS^7,8,22,44^.

The new molecular markers of we have identified unlock unprecedented genetic access to motor neuron subtypes of the spinal cord. Defining the transcriptomes of distinct motor neuron types introduces the possibility of engineering more refined stem cell-derived models and provides a single-cell framework for characterizing the behavior of these rare cell populations in disease.

## Figure Legends

**Extended Data Figure 1:**
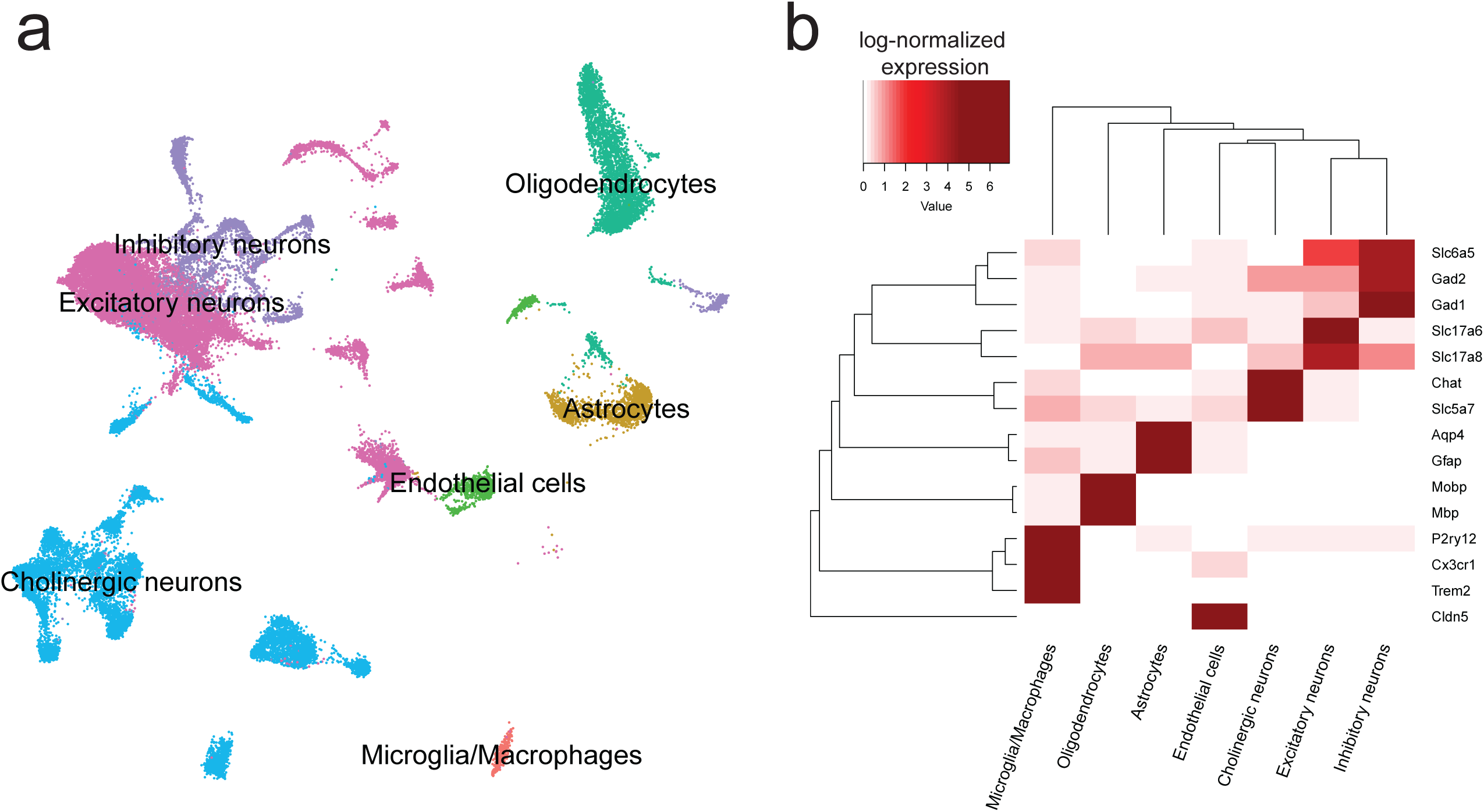
Single-nucleus transcriptional analysis of the adult mouse spinal cord reveals canonical cell types. **a**, Canonical cell class labels, visualized on UMAP. **b**, Average log-normalized marker gene expression across canonical cell classes.

**Extended Data Figure 2:**
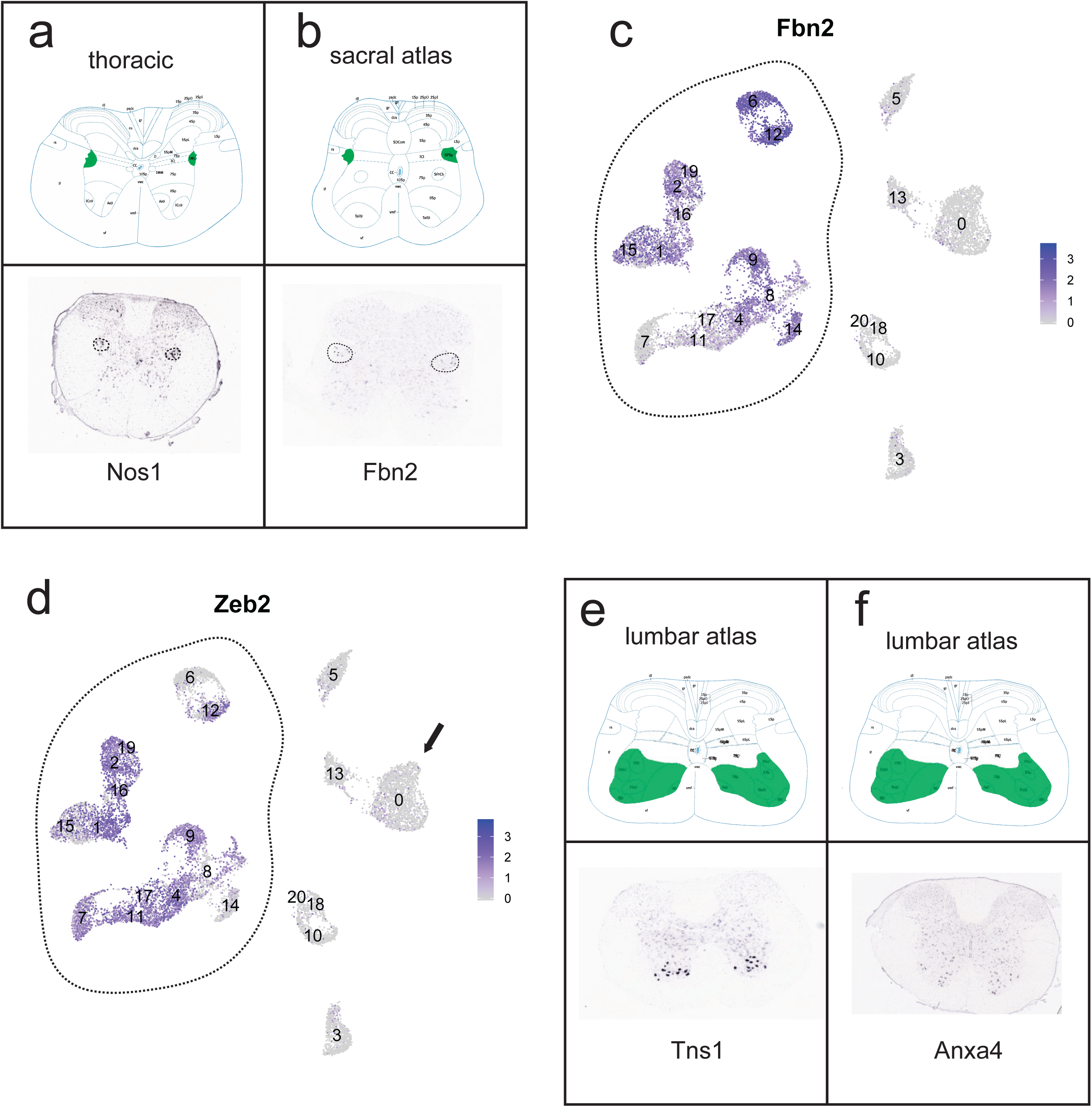
Novel markers of skeletal and visceral motor neurons confirmed by *in situ* hybridization. **a**, Transverse schematic illustrating expected position of visceral motor neurons in lateral autonomic column (LAC, green) in thoracic spinal cord. Below, corresponding *in situ* hybridization against *Nos1*. Dotted circles highlight cell bodies in LAC. **b**, Transverse schematic illustrating expected position of visceral motor neurons in LAC (green) in sacral spinal cord. Below, corresponding *in situ* hybridization against *Fbn2*. Dotted circles highlight cell bodies in LAC. **c**, Average log-normalized expression of *Fbn2* across all cholinergic clusters (labeled), overlaid on UMAP. Dotted line surrounds clusters corresponding to visceral motor neurons. **d**, Average log-normalized expression of *Zeb2* across all cholinergic clusters (labeled), overlaid on UMAP. Dotted line surrounds clusters corresponding to visceral motor neurons, arrow points to α motor neuron cluster (very low *Zeb2* expression). **e**, Transverse schematic illustrating expected position of skeletal motor neurons in ventral horn (VH, green) in lumbar spinal cord. Below, corresponding *in situ* hybridization against *Tns1*, with specific expression in VH. **f**, Transverse schematic illustrating expected position of skeletal motor neurons in ventral horn (VH, green) in lumbar spinal cord. Below, corresponding *in situ* hybridization against *Anxa4*, with specific expression in VH. *In situ* images and transverse schematics adapted from Allen Spinal Cord Atlas^17^.

**Extended Data Figure 3:**
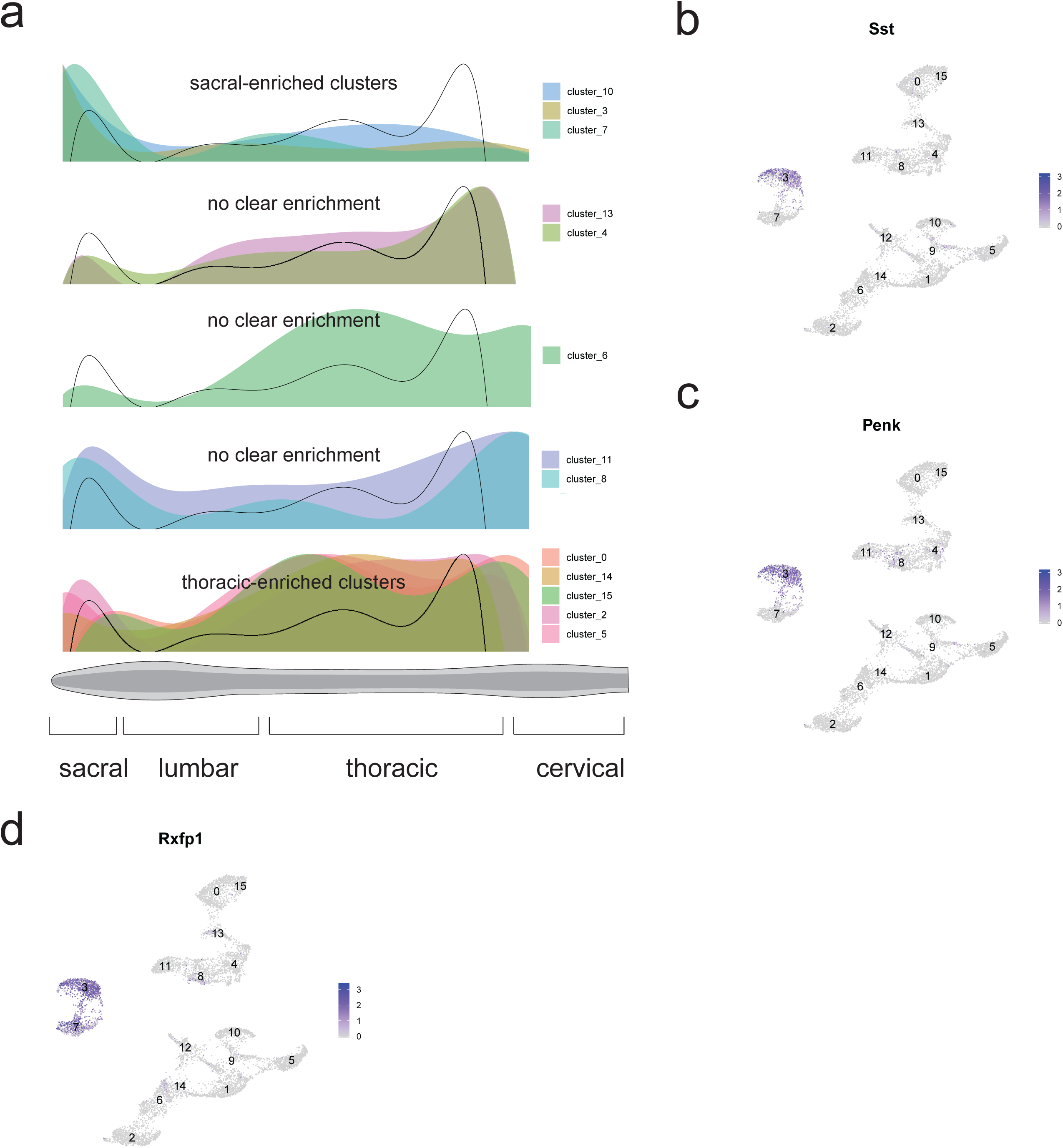
Visceral motor neuron subclusters express selective repertoires of neuropeptides and are spatially distinct. **a**, Transverse schematic illustrating expected position of visceral motor neurons in lateral autonomic column (LAC, green) in thoracic spinal cord. Below, corresponding *in situ* hybridization against *Nos1*. Dotted circles highlight cell bodies in LAC. **b**, Transverse schematic illustrating expected position of visceral motor neurons in LAC (green) in sacral spinal cord. Below, corresponding *in situ* hybridization against *Fbn2*. Dotted circles highlight cell bodies in LAC. **c**, Average log-normalized expression of *Fbn2* across all cholinergic clusters (labeled), overlaid on UMAP. Dotted line surrounds clusters corresponding to visceral motor neurons. **d**, Average log-normalized expression of *Zeb2* across all cholinergic clusters (labeled), overlaid on UMAP. Dotted line surrounds clusters corresponding to visceral motor neurons, arrow points to α motor neuron cluster (very low *Zeb2* expression). **e**, Transverse schematic illustrating expected position of skeletal motor neurons in ventral horn (VH, green) in lumbar spinal cord. Below, corresponding *in situ* hybridization against *Tns1*, with specific expression in VH. **f**, Transverse schematic illustrating expected position of skeletal motor neurons in ventral horn (VH, green) in lumbar spinal cord. Below, corresponding *in situ* hybridization against *Anxa4*, with specific expression in VH. *In situ* images and transverse schematics adapted from Allen Spinal Cord Atlas^17^.

**Extended Data Figure 4:**
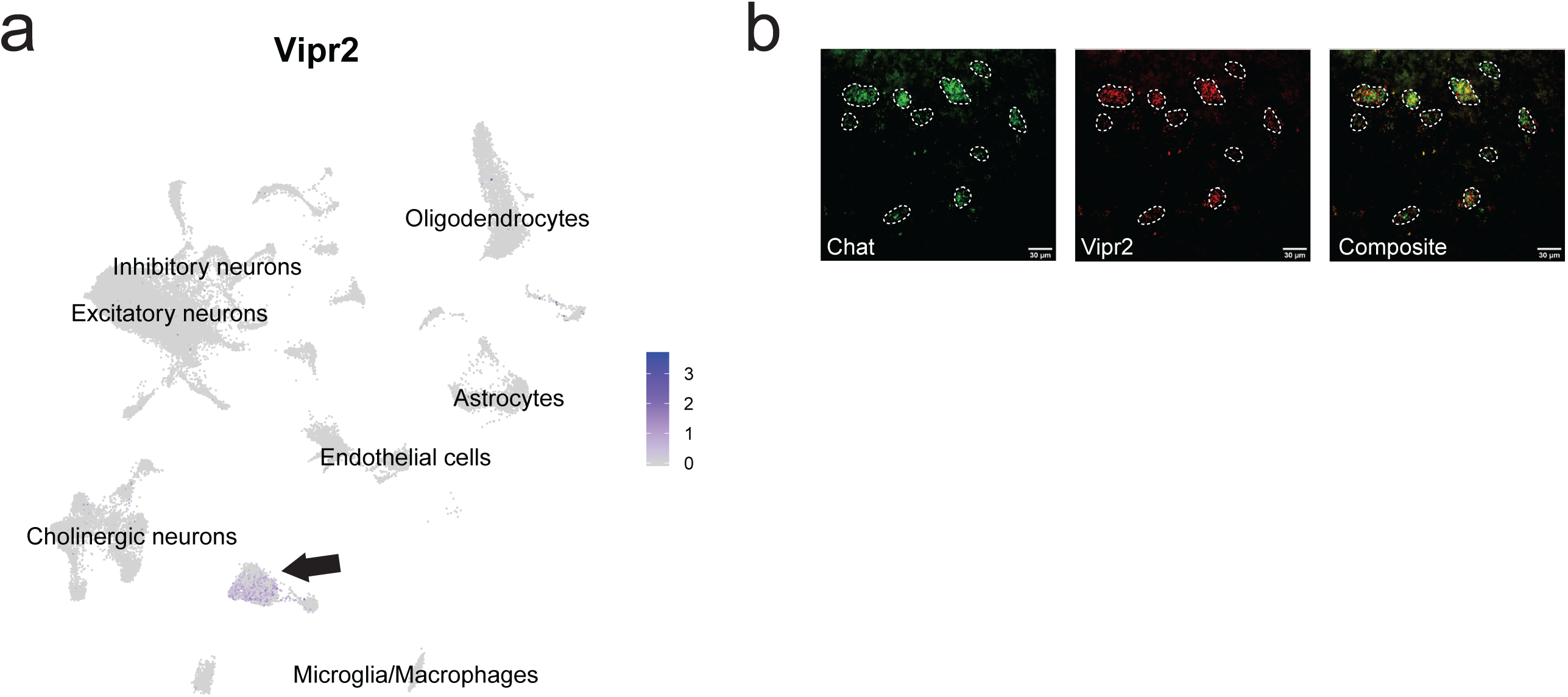
*Vipr2* is a novel, robust, and specific marker of α motor neurons in the spinal cord. **a**, Average expression of *Vipr2* across all spinal cord cell populations (labeled), overlaid on UMAP. Arrow points to α motor neurons cell cluster that expresses *Vipr2.* **b**, *In situ* hybridization of *Chat* and *Vipr2* demonstrates overlap. Image from ventral spinal cord transverse section. Dotted lines mask *Chat*+ cells, scale bars=30 µm.

**Extended Data Figure 5:**
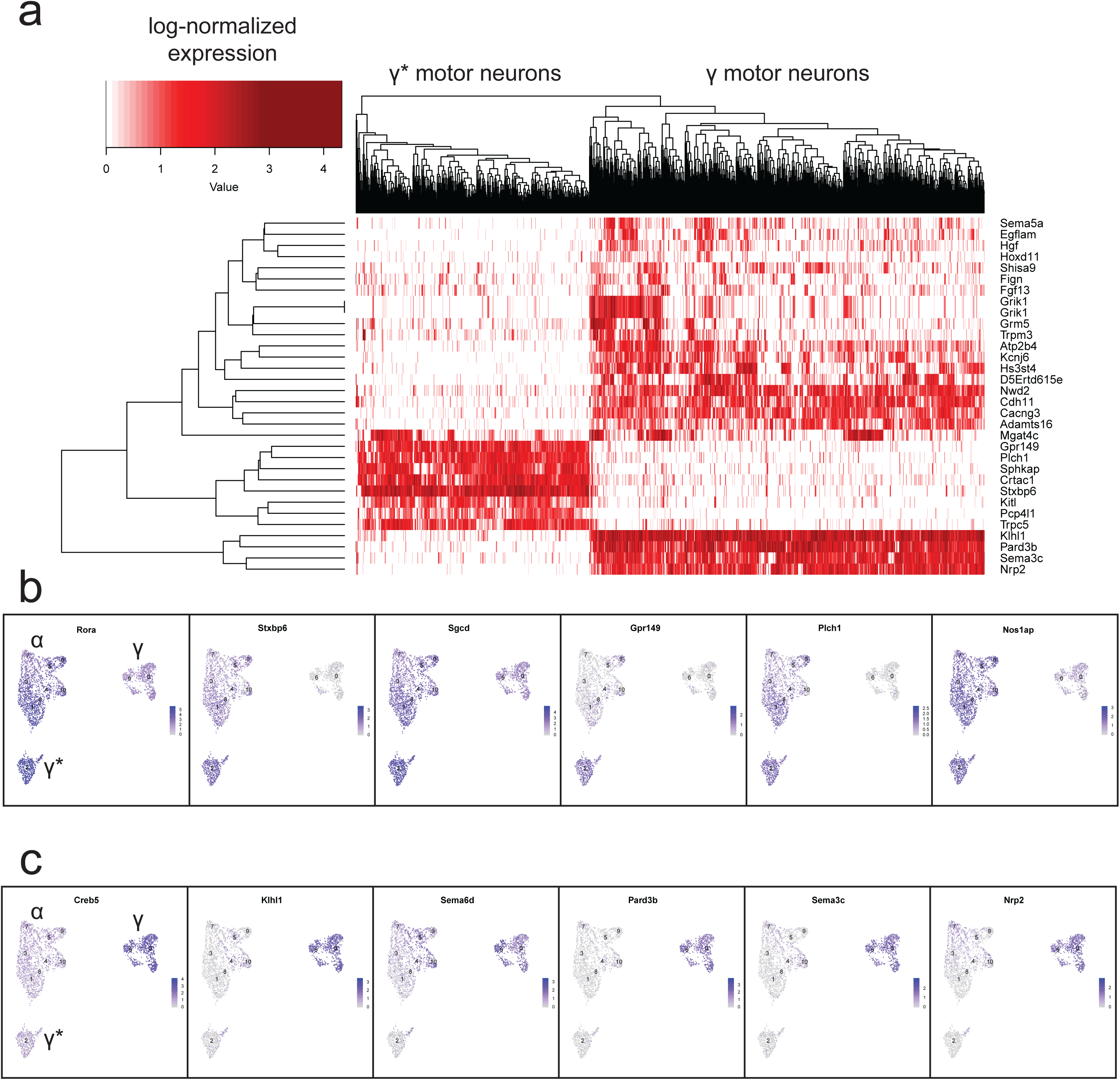
Hierarchical clustering underlines fundamental subdivision within γ motor neurons. **a**, Heatmap with all γ and γ* motor neurons, hierarchically clustered by expression of genes differentially expressed among graph-based clusters. **b**, Average log-normalized expression of genes enriched in γ* motor neurons over γ overlaid on UMAP. α, γ, and γ* populations are labeled. **c**, Average log-normalized expression of genes enriched in γ motor neurons over γ* overlaid on UMAP. α, γ, and γ* populations are labeled. Differentially expressed genes determined by DESeq2 implementation in Seurat (p_adj<0.01, log2-fold change >0.5).

**Extended Data Figure 6:**
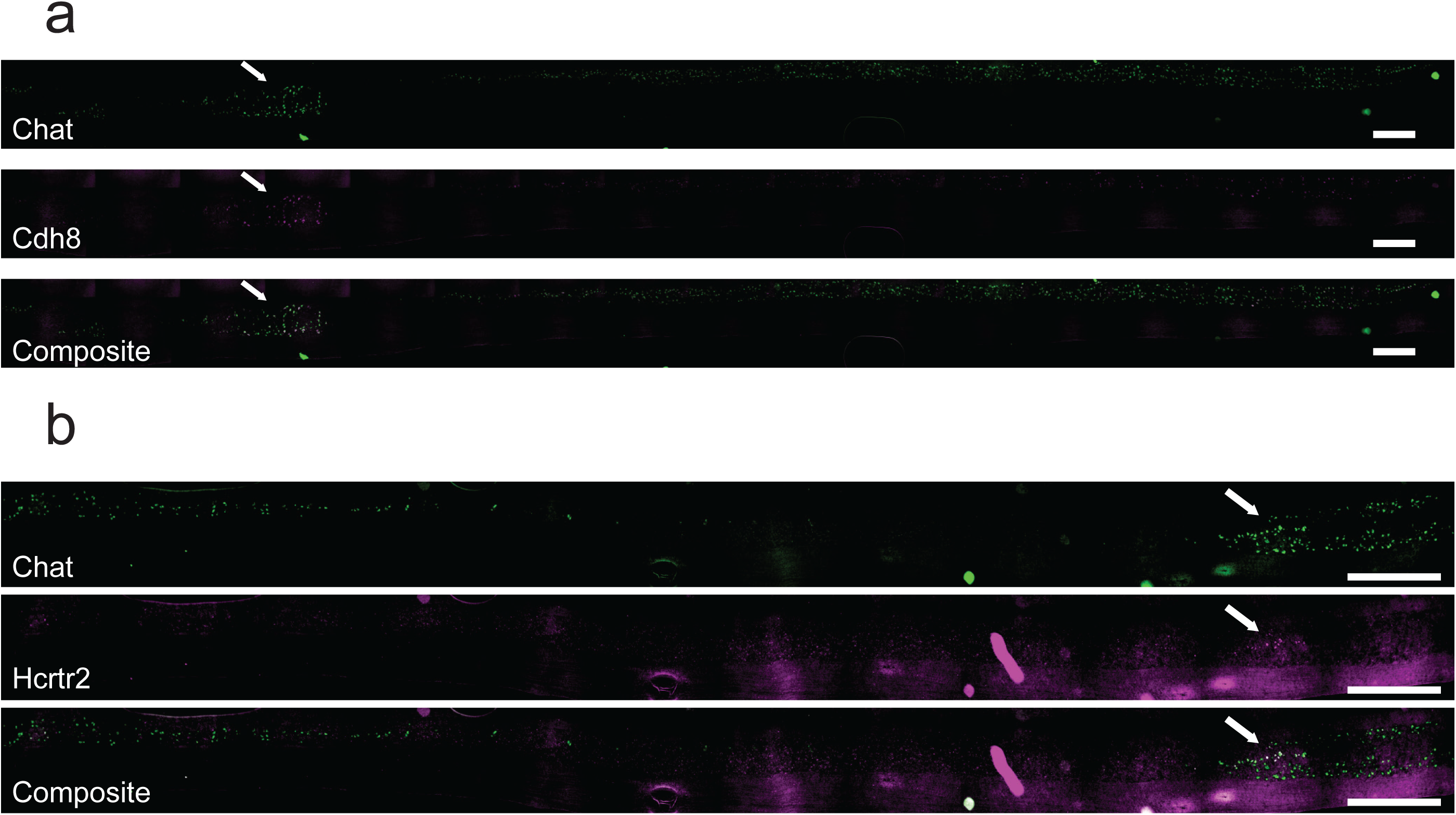
Sagittal *in situ* hybridizations reveals specific localization pattern of α motor neuron subpopulations. **a**, *In situ* hybridization against *Chat* (green) and *Cdh8* (magenta) on sagittal spinal cord section. White arrows indicate position of *Cdh8*-enriched population of motor neurons. **b**, *In situ* hybridization against *Chat* (green) and *Hcrtr2* (magenta) on sagittal spinal cord section. White arrows indicate position of *Hcrtr2*-enriched population of motor neurons. Scale bars=100 µm.

## Acknowledgments

This work was supported by NIH grants R35NS097263 (A.D.G.), R01NS083998 (J.A.K.), P50HG007735 (W.J.G.), the Robert Packard Center for ALS Research at Johns Hopkins (A.D.G.), the Blavatnik Family Foundation (J.A.B.), the Brain Rejuvenation Project of the Wu Tsai Neurosciences Institute (A.D.G.), and the Chan-Zuckerberg Initiative (W.J.G.). Sorting was performed on an instrument in the Shared FACS Facility obtained using NIH S10 Shared Instrument Grant S10RR025518-01. This work used the Stanford Neuroscience Microscopy Service, supported by the grant award NIH NS069375.

## Author Contributions

J.A.B. and A.D.G. designed the experiments and wrote the paper. All authors reviewed and edited the manuscript. J.A.B. performed experiments and computational analysis of the data. S.K. helped plan and perform experiments and provided advice on analysis. L.N. helped with mouse husbandry. K.A.G., A.K., P.T.H., helped perform experiments. J.L.S. and J.A.K. provided advice on analysis and helped plan experiments. W.J.G. helped analyze data and provided advice on designing experiments.

## Methods

### Data reporting

No statistical methods were used to predetermine sample size.

### Data availability

All sequencing data will be deposited in NCBI-GEO prior to publication.

### Code availability

All code used to analyze data and generate figures is available on request from corresponding author.

### Mouse crosses

CHaT-IRES-Cre (Chat-CRE/Chat-CRE) mice were purchased from JAX (stock no. 006410m B6;129S6-*Chat*^*tm2(cre)Lowl*^/J) and crossed with ROSA^nT-nG^ /ROSA^nT-nG^ (stock no. 023035, B6;129S6-*Gt(ROSA)26Sor*^*tm1(CAG-tdTomato*, -EGFP*)Ees*^/J). F1 heterozygous reporter mice were aged to P100-150, and then sacrificed for subsequent sequencing and *in situ* experiments.

### Mouse nuclei collection

Four to 8 mice in five independent experiments were euthanized with CO2 and decapitated caudal to the brain stem. Their spinal columns were severed just caudal to the sacral spinal cord, and rapidly cut out as described^11^. Briefly, a blunt 18-gauge syringe containing ice-cold PBS was inserted into the caudal end of the spinal cord and used for rapid hydraulic extrusion of the entire, intact cord. Two spinal cords at a time were homogenized with a Dounce Homogenizer in 2 mL of nuclei extraction buffer (neB, Supp. Methods Table 1). Spinal cords were homogenized with 10 strokes of Pestle A, followed by 5 strokes of Pestle B. The entire homogenate was transferred to a 25 mL, round-bottom plastic ultracentrifuge tube and 8 mL of nuclei spin buffer 1 (nsB1, Supp. Methods Table 1) was added. Five mL of nuclei spin buffer 2 (nsB2, Supp. Methods Table 1) were layered gently underneath the homogenate, and the gradient was spun for 15 minutes at 4000* g in a 4°C, benchtop swinging bucket centrifuge (Beckman 5810R). The supernatant was rapidly discarded, and the nuclei were gently resuspended in 5 mL nsB1. Five mL of nuclei spin buffer 3 (nsB3) were gently layered underneath, and the resulting gradient was spun for 15 minutes at 4000* g at 4° C. The supernatant was rapidly discarded, and the pellet was resuspended in 400 uL nuclei FACS buffer (nfB). Next, 0.5 uL DAPI was added to enable doublet discrimination on the sorter.

**Table 1:**
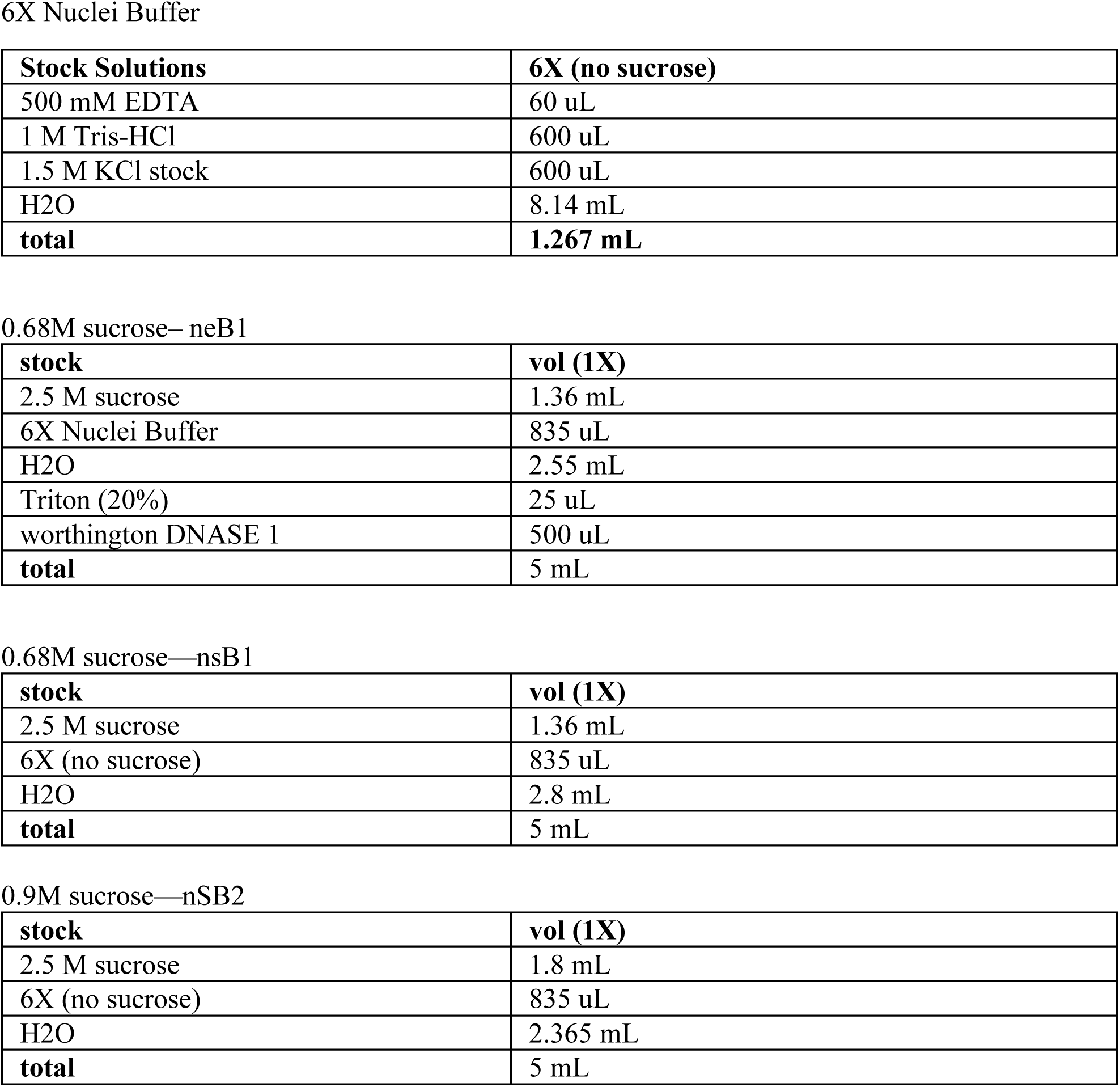
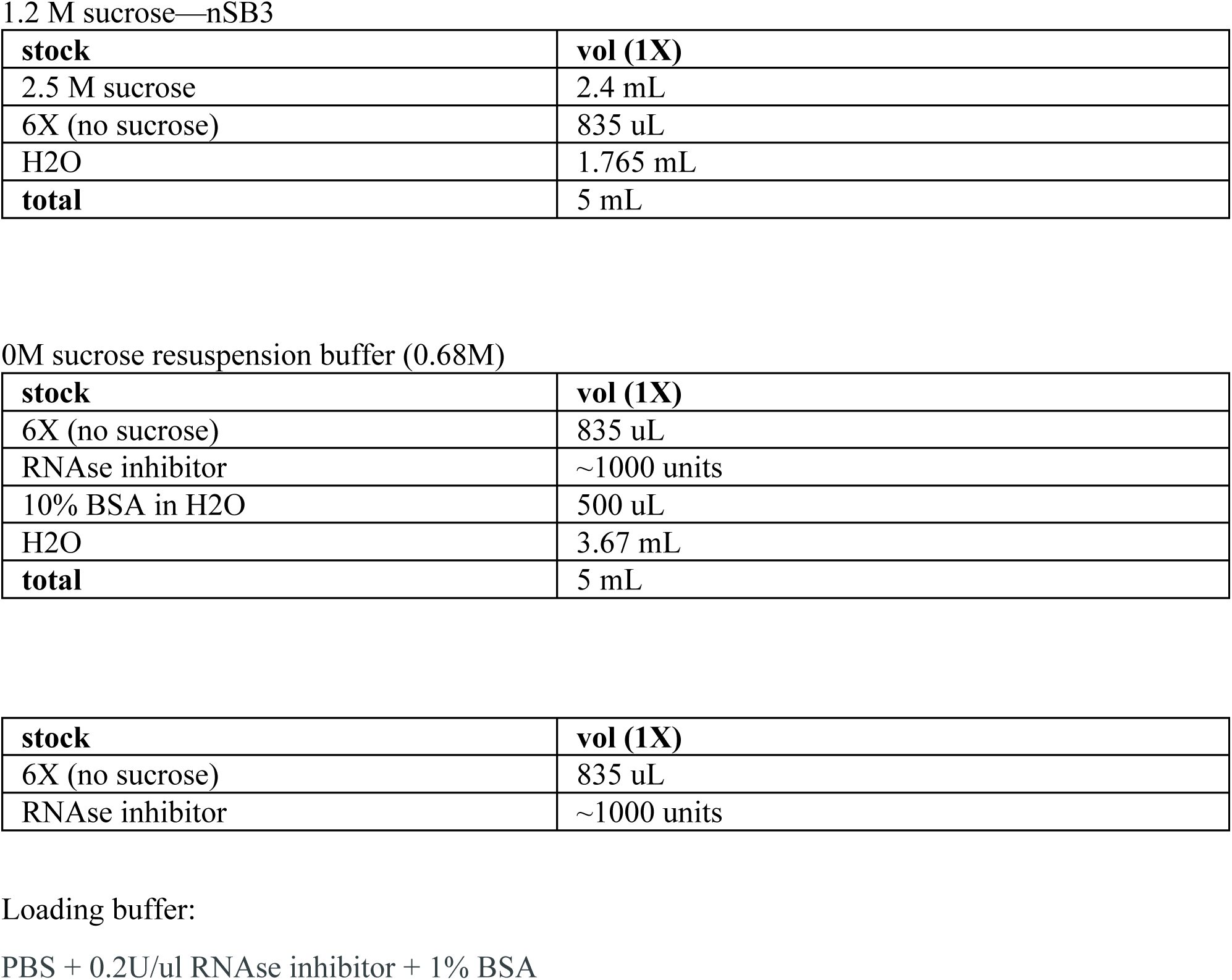
Buffers.

### Fluorescence activated nuclei sorting (FANS)

FANS was performed as described on a BD Biosciences FACSAria II flow cytometer. Briefly, following calibration of the FACS machine or power-up of the flow cytometer, single nuclei were gated using forward scatter (FSC), side-scatter (SSC), and DAPI measurements to ensure that doublets were gated out. Following this initial gating, EGFP+/tdTomato-nuclei were identified using a 2-dimensional scatterplot. These nuclei stood out from the main population, enabling double-gating. To ensure that our samples would contain both cholinergic and non-cholinergic nuclei, we then sorted ∼10,000-15,000 EGFP+/TdTomato-nuclei and 20,000-25,000 EGFP-/TdTomato+ nuclei into 400 uL 10X loading buffer (see below). After sorting was completed, this nuclei mixture was spun at 400xg for 5 minutes and resuspended in 10X loading buffer at 1000 cell/ul estimated under the assumption that only 50% of cells sorted by flow cytometry are recovered.

### Droplet-based snRNA-seq

For droplet-based snRNA-seq, libraries were prepared using the Chromium Single Cell 3′ Reagent Kits v.3 according to the manufacturer’s protocol (10x Genomics). For all experiments 6000-8000 cells were targeted for each sample. The generated snRNA-seq libraries were sequenced using NextSeq 500/550 High Output v2 kits (150 cycles) and NovaSeq 6000 S4 (150 cycles) to an average read depth of ∼100,000 reads/cell.

### Mouse CNS *in situ* hybridization

Mice were humanely euthanized using CO2 for 5 minutes prior to decapitation. The spinal cord was rapidly hydraulically extruded, as above, and briefly dried. For longitudinal sections, the spinal cord was placed into a plastic freezing mold and frozen briefly on dry ice, before adding

O.C.T Compound freezing media (Tissue-Tek) to ensure that the spinal cord was frozen completely flat to the mold. For transverse sections, spinal cords were cut into three segments— roughly lumbosacral, thoracic, and cervical. Sections were mounted and cut to a thickness of 20 um using a Leica CM3050 S Cryostat. Sections were processed immediately or stored for 1-2 weeks at -80° C.

Sections were processed for RNAScope (ACD Biosciences) according to manufacturer instructions for fresh frozen tissue. All *in situ* hybridization probes used are listed in Supp. Methods Table 2. Briefly, tissue sections were fixed with 4% PFA at 4° C, and subsequently dehydrated with 5 minutes each of 50% EtOH, 70% EtOH, and finally 100% EtOH. Samples were dried and a hydrophobic barrier was drawn around each section. Samples were then incubated with Protease IV (ACD Biosciences) for 15 minutes at room temperature before removing and washing in 1X PBS. Before staining, probes were equilibrated to 40° C for 10 minutes and then cooled to room temperature. Mixtures of probes in channels 1-3 were diluted, and then incubated on samples for 2 hrs in a humidified 40° hybridization oven. Samples were washed twice in PBS, then incubated in the hybridization oven at 40° with Amp-1-FL, Amp-2-FL, Amp-3-FL, and Amp-4-FL for 30, 15, 30, and 15 minutes respectively (with 2X washes in between each incubation). Samples were then washed twice, briefly dried, and mounted with DAPI-containing VectaShield mounting media (Vector Laboratories, H-1200-10). RNAscope *in situ* hybridization images were taken using a Zeiss Axio Imager M1 and Axiovision software or a Zeiss 710 inverted confocal microscope and Zen software. Image analysis was performed using ImageJ.

**Table 2:** RNAscope *in situ* probe list.

| **Probe target** | **Channel** | **Catalog Number** |
| --- | --- | --- |
| Mm-Chat | C1,C2 | 408731-C1,C2 |
| Mm-Prkcd | C1 | 441791 |
| Mm-Vipr2 | C1 | 465391 |
| Mm-Piezo2 | C3 | 400191-C3 |
| Mm-Htr1d | C1, C2 | 315871-C1,C2 |
| Mm-Kcnq5 | C3 | 511131 |
| Mm-Pard3b | C1 | custom |
| Mm-Cdh8 | C1 | 485461 |
| Mm-Stxbp6 | C3 | 443281-C3 |

### Analysis of snRNA-seq data

Counts per gene were generated by aligning reads to the mm10 genome (GrCm38—mm10 (GCF_000001635.20) using CellRanger software (v.3.0.0) (10x Genomics) running on the Sherlock Stanford Computing Cluster. To account for unspliced nuclear transcripts, reads mapping to pre-mRNA were counted. After this, the CellRanger aggr pipeline was used to aggregate all libraries and normalize the read depth between libraries before data merging (with the default parameters) to generate a gene-count matrix. Doublets and empty droplets were excluded from downstream analysis using cell-detection method provided by CellRanger (10X genomics). One doublet-containing cluster was subsequently removed.

### Clustering and normalization

Single-nucleus transcriptome normalization, batch correction, clustering and dimensionality reduction were performed performed using the Seurat R Package^70^ (https://satijalab.org, V3.0). All 43,890 transcriptomes were normalized for read-depth with the CellRanger aggr function, and then loaded into Seurat. The nuclei were batch-corrected using the Seurat *Integrate* function as previously described^70^, which enables cells from different experiments to be projected into the same high and low-dimensional spaces. Principal component analysis (PCA) was performed on the whole dataset, and the top 15 components were used to generate a uniform manifold approximation (UMAP)^71^. The same principal components were used to perform graph-based clustering via the *FindClusters* function, which identified 39 total clusters.

#### Differential expression and subcluster identification

To annotate clusters, a manually curated list of markers for major cell types was assembled from the literature – including excitatory interneurons, inhibitory interneurons, cholinergic neurons, oligodendrocytes, endothelial cells, astrocytes, and microglia. Average expression of these marker genes was calculated using Seurat (*AverageExpression*). Clusters that were positive for these marker genes were annotated by cell type (e.g. *Aqp4+* clusters were classified as astrocytes). To identify specific marker genes for each neuronal subtype, differential gene expression analysis was performed using the Seurat *FindAllMarkers* function, which leverages DESeq2 differential expression testing. Briefly, differentially expressed genes were identified for each subtype with respect to all remaining cells. Each subtype was down-sampled to 100 cells in order to compensate for stoichiometric biases. An FDR-corrected p-value threshold was set at p<0.01, and only markers with a log2-fold change>0.5 were included in subsequent analysis. To find robust markers, lists of differentially expressed marker genes were rank order based on the percent expression level in all cells not included in the subtype of interest (low background expression yields high rank). Even though this occasionally resulted in the prioritization of markers that are not ubiquitously expressed in the subtype of interest, we determined this method empirically produces robust markers and view the loss of signal is some cells a likely result of a high dropout rate for single-nucleus sequencing. However, only genes detected in at least 25% of the cells within the given identity class were considered as candidate subtype markers. Skeletal and visceral motor neurons, as well as cholinergic interneurons, were each separately subsetted, subclustered, and umap-projected. Differential expression analysis was performed on clusters within each population, as above and explained in the main text.

### Visceral motor neuron spatial localization

We determined where cells within each visceral motor neuron subcluster localize along this axis by the following method. Briefly, a list of genes that localize to the intermediolateral cell column (iMLC) was downloaded from the Allen Spinal Cord Atlas^17^ (Supp. Methods Table 3). Differential expression analysis among visceral subclusters was performed *exclusively for* this list of genes, such that we were able to identify 29 enriched genes (marker genes) within this ∼100-gene list across visceral motor neuron subclusters. We then counted the number of iMLC cells that were positive for each gene in all available adult spinal cord sections in the Allen Spinal Cord Atlas and noted the relative position of each positive cell along the rostral-caudal spinal cord axis.

**Table 3:**
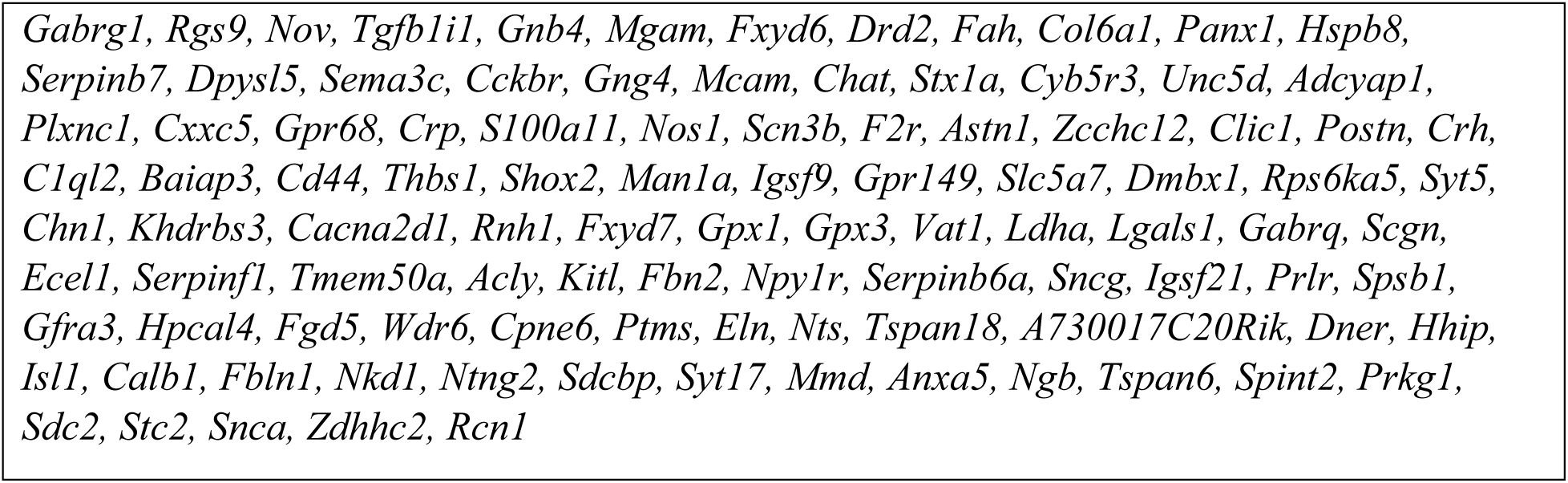
iMLC-expressed gene list (Allen Spinal Cord Atlas^17^)

For each marker gene, we normalized the number of positive cells at each rostral-caudal position to the mean positive cell count for that gene across all sections. We then divided each point by the maximum number of cells in any section that were positive for that gene to determine the *relative* enrichment in different sections of the spinal cord. Owing to the sparsity of the data, we fit a polynomial function (order=8) to the normalized positive cell counts, producing 29 independent polynomial functions that reflect the density of cells expressing each gene over the adult mouse spinal cord. We then calculated the average expression of each gene across all clusters, and for each gene we calculated the log2-fold change of each cluster compared to the cells in the other populations. If the log2-FC>0 and the FDR-adjusted p-value was <0.01, we considered a cluster enriched for expression of that gene. Otherwise, we did not. For each cluster, we calculated a weighted positional density by multiplying the log2-FC of each enriched genes by its polynomial density, summing over all enriched genes, and renormalizing the cumulative density. This effectively generates an estimated positional density for each cluster along the rostral-caudal spinal cord spatial atlas as reported.

## Supplemental Methods

